# Association of OXA-1 and TEM-1 Genes with Antibiotic Resistance to Piperacillin/Tazobactam in ESBL-Producing Pathogens: Insights from a Multi-Center Analysis

**DOI:** 10.1101/2025.04.18.649505

**Authors:** Edwin Kamau, Brendan M. Wong, John L MacArthur, Jamie L Dombach

**Affiliations:** Department of Pathology and Area Laboratory Services, Tripler Army Medical Center, Honolulu, HI, USA; Adult Intensive Care Unit, Tripler Army Medical Center, Honolulu, HI, USA; Department of Clinical Investigation, Tripler Army Medical Center, Honolulu, HI, USA

## Abstract

**Background:** The emergence of extended-spectrum β-lactamase (ESBL)-producing *Escherichia coli* and *Klebsiella pneumoniae* presents significant challenges in treating infections caused by these pathogens. This multi-center retrospective study investigated the prevalence of OXA-1 and TEM-1 genes in ESBL-producing *E. coli* and *K. pneumoniae*, along with their association with piperacillin/tazobactam susceptibility.

**Methods:** Clinical isolates were collected from three institutions as part of routine patient care: Tripler Army Medical Center (TAMC) in Hawaii, Madigan Army Medical Center (MAMC) in Washington, and Brooke Army Medical Center (BAMC) in Southern Texas. A total of 416 isolates were analyzed through genome sequencing and CLSI-guided susceptibility testing.

**Results:** OXA-1 and TEM-1 β-lactamase enzymes were present in 20.9% (73/349) and 38.7% (135/349) of the *E. coli* isolates, respectively. Relative risk analysis of non-susceptibility to piperacillin/tazobactam across isolates from the three study sites revealed a highly significant association for OXA-1 (*P* < 0.001), whereas no significant associations were observed for TEM-1 (*P* = 0.424) or the combination of OXA-1 and TEM-1 (*P* = 0.082). When analyzed by institution, the relative risk of non-susceptibility to piperacillin/tazobactam remained highly significant for OXA-1 at TAMC and MAMC (*P* < 0.001 for both) but was not significant at BAMC (*P* = 0.21). OXA-1 and TEM-1-positive variants showed a significant association with genes conferring resistance to other antibiotics.

**Conclusions:** The OXA-1 gene plays a key role in resistance to piperacillin/tazobactam in ESBL-producing organisms, with geographic differences in non-susceptibility observed. Genetic profiling and localized data are crucial for optimizing antibiotic therapy and improving treatment outcomes.

## INTRODUCTION

Extended-spectrum β-lactamase (ESBL)-producing Gram-negative pathogens have emerged as a significant public health threat since their discovery in the early 1980s. These pathogens have developed resistance to expanded-spectrum β-lactam antibiotics, posing challenges in both hospital-associated (HA) and community-acquired (CA) infections (1,2). The global spread of ESBL-producing Enterobacterales has necessitated the exploration of alternative treatment options to combat these resistant strains (1).

Piperacillin/tazobactam has been proposed as a potential carbapenem-sparing alternative for managing infections caused by ESBL-producing *Escherichia coli* and other Enterobacterales. However, clinical studies on the efficacy of penicillin/inhibitor combinations against ESBL-producers have yielded contradictory results (3-7). The MERINO trial, a sentinel study investigating bacteremia due to ESBL strains, reported a 30-day mortality rate of 12.3% for patients treated with piperacillin/tazobactam compared to 3.7% for those treated with meropenem (*P* = 0.002) (6). Post-hoc analysis of the MERINO trial suggested that narrow-spectrum β-lactamases played a significant role in the poor performance of piperacillin/tazobactam compared to meropenem (7). Importantly, the clinical outcome for patients receiving piperacillin/tazobactam may depend, at least in part, on the minimum inhibitory concentration (MIC) for this antimicrobial (4).

Since the MERINO trial, studies have demonstrated that several factors contribute to the variable resistance observed in ESBL-producers when treated with penicillin/inhibitor combinations. These factors include the production of multiple β-lactamases, such as OXA-1 and TEM-1, which are poorly inhibited by these combinations (8-10). Additionally, hyperproduction of target β-lactamases and bacterial impermeability further complicate treatment outcomes. A study conducted by Livermore et al. assessing ESBL-producing isolates from the United Kingdom found that the presence of OXA-1 but not TEM-1 was strongly associated with reduced susceptibility to piperacillin/tazobactam and amoxicillin/clavulanate (8), highlighting the enzyme’s role in resistance mechanisms. Similar findings were demonstrated in ESBL-producing isolates from Canada (10). A study conducted by Rodríguez-Villodres demonstrated that piperacillin/tazobactam was less effective against ESBL *E. coli* carrying *bla*^TEM^ due to increase in copy numbers and transcription levels of the gene, and patients were at high risk of therapeutic failure (11). These findings underscore the importance of considering the presence of OXA-1 and TEM-1 when evaluating the efficacy of penicillin/inhibitor combinations against ESBL-producers.

As in most settings, piperacillin/tazobactam is one of the most widely prescribed empiric treatment options in military treatment facilities (MTFs) within the Defense Health Agency (DHA). In this study, we performed a multi-site retrospective study to investigate the prevalence of OXA-1 and TEM-1 in ESBL-producing *E. coli* and *Klebsiella pneumoniae* in our patient populations. We explored the association between the minimal inhibitory concentrations (MICs) of piperacillin/tazobactam and the presence of OXA-1 and/or TEM-1 carrying ESBL isolates.

## RESULTS

A total of 416 ESBL-producing organisms, including 349 *E. coli* and 67 *K. pneumoniae*, were analyzed in this study. There were 106, 94, and 216 isolates from Tripler Army Medical Center (TAMC) in Hawaii, Madigan Army Medical Center (MAMC) in Washington, and Brooke Army Medical Center (BAMC) in Southern Texas, respectively, of which 93, 85, and 171 were *E. coli*. There were 343 *E. coli* isolates with known sequence types (STs), of which 183 (52.4%) were ST131. Other STs with eight or more isolates were ST1193 (n=26), ST38 (n=16), ST998 (n=14), ST69 (n=10), ST10 (n=10), ST636 (n=9), and ST648 (n=8). Six isolates did not have known STs. When we evaluated the proportion of each ST in each institution, BAMC had the highest proportion of ST131 isolates (57.3% [98/171]), while MAMC had the lowest (47.1% [40/85]). Although the total number of isolates was small, the proportions of ST1193 and ST998 were highest in TAMC and lowest in BAMC, whereas seven of the eight ST648 isolates were found in BAMC and none in TAMC. Most isolates harbored a CTX-M ESBL enzyme, predominantly the CTX-M-15 β-lactamase, which was present in 50.1% (175/349) of the isolates, with 58.9% (103/175) being ST131. The narrow-spectrum OXA-1 and TEM-1 β-lactamase enzymes were present in 20.9% (73/349) and 38.7% (135/349) of the *E. coli* isolates, respectively. Of the CTX-M-15 ESBL variants, 41.7% (73/175) were OXA-1 positive, and 34.3% (60/175) were TEM-1 positive. Of the CTX-M-15/ST131 isolates, 54.4% (56/103) were OXA-1 positive, and 36.9% (38/103) were TEM-1 positive; OXA-1 was only detected in the CTX-M-15 ESBL type variant. Other notable β-lactamase variants identified included CTX-M-27 (n=98, 28.1%), CTX-M-14 (n=36, 10.3%), CTX-M-55 (n=24, 6.9%), and CTX-M-3 (n=8, 2.3%). For *K. pneumoniae*, 92.5% (62/67) had known STs, with the two predominant STs being ST307 (n=9) and ST45 (n=8).

The highest proportion of isolates carrying CTX-M-15 was found in BAMC (59.1% [101/171]), while the lowest was in TAMC (36.6% [34/93]). We evaluated whether there was a significant difference in the proportion of CTX-M-15 variants between institutions and determined there was a significant difference (χ^2^=12.63; *P*=0.0018). Pairwise comparison for the proportion test revealed no difference in the CTX-M-15 variant between TAMC and MAMC (*P* = 0.156), and MAMC and BAMC (*P* = 0.069). However, there was a difference between TAMC and BAMC (*P* < 0.001), revealing that the significant difference was primarily between TAMC and BAMC. For the CTX-M-27 ESBL variant type, TAMC had the highest proportion of isolates (38.7% [34/93]), whereas the lowest proportion was in BAMC (24.0% [41/171]). Analysis revealed a statistically significant association between the institutions and the distribution of the CTX-M-27 variant (*P* = 0.029), specifically between TAMC and MAMC (*P* = 0.045), and TAMC and BAMC (*P* = 0.012), but not between MAMC and BAMC (*P* = 0.898). This indicated that the significant differences were primarily between TAMC and both MAMC and BAMC. For the CTX-M-14 ESBL variant type, TAMC (15.0% [14/93]) had the highest proportion, while the lowest was in BAMC (5.8% [10/171]), suggesting that the distribution of this variant was significantly associated with the institution (χ^2^=7.27; *P*= 0.026). Pairwise comparison for the proportion test revealed a significant difference in the proportion of the CTX-M-14 variant between TAMC and BAMC (*P* = 0.013), and MAMC and BAMC (*P* = 0.026), but not between TAMC and MAMC (*P* = 0.860).

Using the current Clinical and Laboratory Standards Institute (CLSI) susceptibility breakpoint of ≤8 mg/L for Enterobacterales, we examined the association between the presence of the OXA-1 or TEM-1 genes and susceptibility to piperacillin/tazobactam. MIC data were available for 88.8% (310/349) of the isolates, of which 92.3% (286/310) were susceptible to piperacillin/tazobactam. Specifically, 95.6% (237/248) of OXA-1 negative isolates were susceptible to piperacillin/tazobactam, compared to 83.9% (52/62) of OXA-1 positive isolates. Figure 1A illustrates the number of isolates from each institution and their susceptibility to piperacillin/tazobactam based on the presence or absence of the OXA-1 or TEM-1 genes. Figure 1B depicts the percent susceptibility based on the presence or absence of these genes. Notably, OXA-1 negative isolates exhibited significantly higher susceptibility in TAMC (96.3% vs. 62.5% OXA-1 positive) and MAMC (98.5% vs. 64.7% OXA-1 positive) compared to BAMC, where OXA-1 negative isolates had lower susceptibility (93.1%) than OXA-1 positive isolates (97.1%).

**Figure 1.**
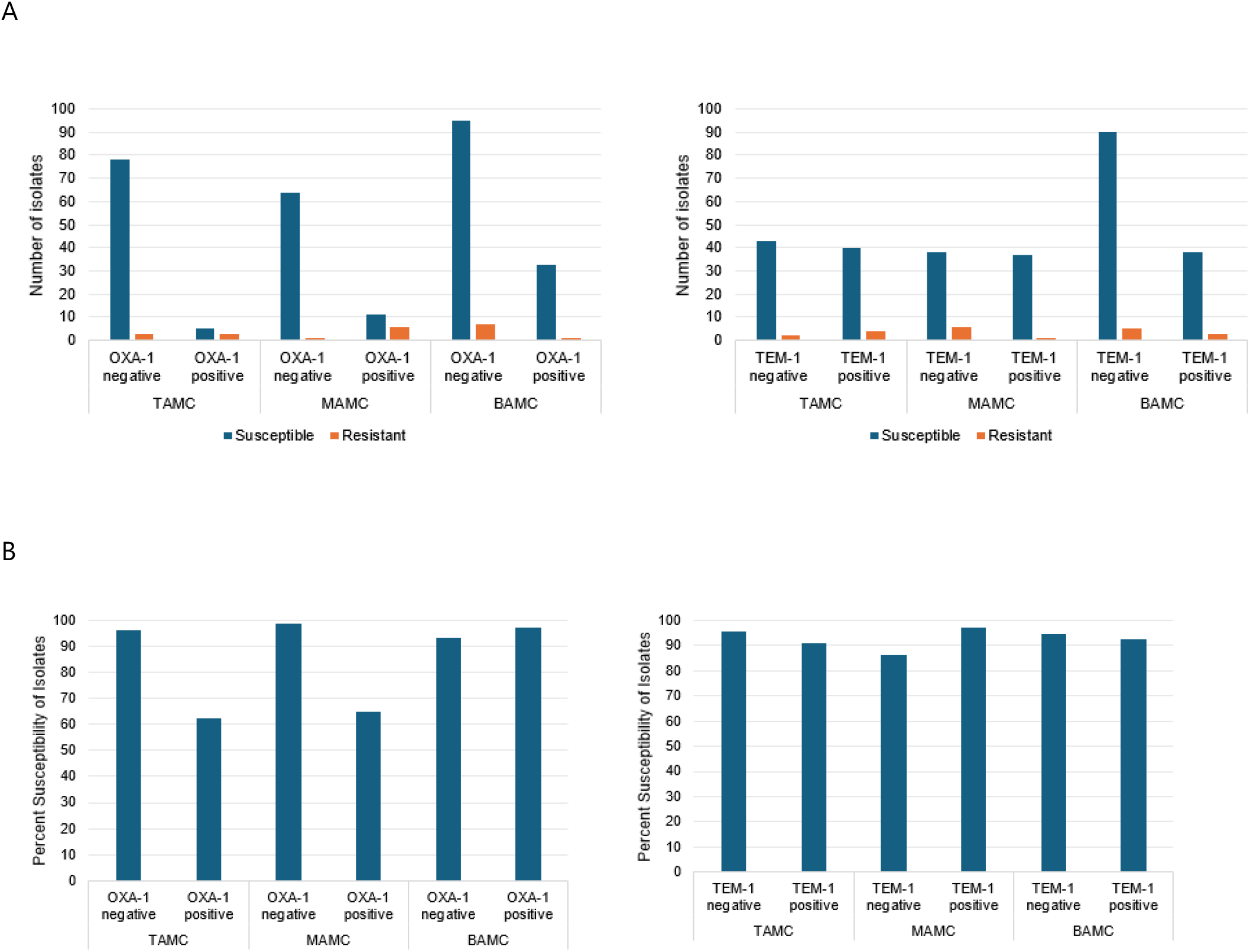
Plot showing the number of *E. coli* isolates present in each institution (A) and percent susceptible isolates (B).

Next, we evaluated the association between the presence of the OXA-1 gene and susceptibility to piperacillin/tazobactam in all ESBL-producing isolates in the three study sites and found a significant association between the presence of OXA-1 and susceptibility to piperacillin/tazobactam (χ^2^= 6.99; *P* = 0.0082). Notably, all OXA-1 isolates were CTX-M-15, representing 41.7% (73/175) of this variant type. We found a significant difference in the proportions of susceptibility to piperacillin/tazobactam between OXA-1 positive and OXA-1 negative *E. coli* bacteria *(P=* 0.0074).

We analyzed the data by institution for the association between the presence of the OXA-1 gene and susceptibility to piperacillin/tazobactam across the three institutions (Table 1). Data from TAMC and MAMC demonstrated a statistically significant association, whereas no such association was observed at BAMC. When analyzing the CTX-M-15/ST131 isolates from the three institutions combined, 54.4% (56/103) were OXA-1 positive. However, there was no statistically significant association between the presence of the OXA-1 gene and susceptibility to piperacillin/tazobactam (χ^2^ = 0.113, *P* = 0.737; proportion test statistic = 0.798, *P* = 0.479). Furthermore, none of the isolates carrying CTX-M-27, CTX-M-14, or CTX-M-55 ESBL enzyme types were OXA-1 positive.

**Table 1:** Data showing chi-square tests for independence and proportion tests across the three study sites.

| institution | category | chi-square | P | Z-statistic | p-value (proportion) |
| --- | --- | --- | --- | --- | --- |
| TAMC | Susceptible | 6.72 | **0.0095** | 2.55 | **0.0107** |
| MAMC | Susceptible | 14.29 | **0.0002** | 3.78 | **0.0002** |
| BAMC | Susceptible | 0.92 | 0.337 | 0.96 | 0.337 |

We next assessed the association between the TEM-1 β-lactamase enzyme and susceptibility to piperacillin/tazobactam. Among all ESBL-producing isolates, chi-square tests for independence and proportion tests indicated no statistically significant association between the presence of the TEM-1 gene and susceptibility to piperacillin/tazobactam (data not shown). Among the CTX-M-15 isolates, 34.3% (60/175) were TEM-1 positive. Of these, 150 isolates (55 TEM-1 positive and 95 TEM-1 negative) had minimum inhibitory concentration (MIC) data available for piperacillin/tazobactam. Analysis of this demonstrated no significant association between the presence of the TEM-1 gene and susceptibility to piperacillin/tazobactam (data not shown).

We conducted a relative risk analysis of non-susceptibility to piperacillin/tazobactam across isolates from the three study sites. The analysis revealed a highly significant association for OXA-1 (*P* < 0.001), whereas no significant associations were observed for TEM-1 (*P* = 0.424) or the combination of OXA-1 and TEM-1 (*P* = 0.082) (Table 2). When analyzed by institution, the relative risk of non-susceptibility to piperacillin/tazobactam remained highly significant for OXA-1 at TAMC and MAMC (*P* < 0.001 for both) but was not significant at BAMC (*P* = 0.21). These findings were consistent with the results of chi-square tests for independence and proportion tests. Analyses of TEM-1 and the combination of OXA-1/TEM-1 by individual institution also showed no significant associations (Table 2).

**Table 2:** Relative risk analysis of non-susceptibility to piperacillin/tazobactam.

| All Isolates |  |  |  |  |
| --- | --- | --- | --- | --- |
|  | RR of MIC<br>>8 mg/m: | 95%<br>lower CI | 95% Upper<br>CI | P |
| **OXA-1** | 3.85 | 1.7 | 8.57 | **<0.001** |
| **TEM-1** | 0.92 | 0.39 | 2.16 | 0.424 |
| **OXA-1 + TEM-1** | 2.59 | 0.68 | 9.86 | 0.082 |
| TAMC |  |  |  |  |
|  | RR of MIC<br>>8 mg/m: | 95%<br>lower CI | 95% Upper<br>CI | P |
| **OXA-1** | 10.13 | 2.43 | 42.14 | **<0.001** |
| **TEM-1** | 2.05 | 0.39 | 10.61 | 0.2 |
| **OXA-1 + TEM-1** | 17.6 | 7.51 | 41.23 |  |
| MAMC |  |  |  |  |
|  | RR of MIC<br>>8 mg/m: | 95%<br>lower CI | 95% Upper<br>CI | P |
| **OXA-1** | 22.94 | 2.96 | 177.96 | **<0.001** |
| **TEM-1** | 0.193 | 0.024 | 1.53 | 0.06 |
| **OXA-1 + TEM-1** | 1.54 | 0.211 | 11.25 | 0.33 |
| BAMC |  |  |  |  |
|  | RR of MIC<br>>8 mg/m: | 95%<br>lower CI | 95% Upper<br>CI | P |
| **OXA-1** | 0.43 | 0.055 | 3.36 | 0.21 |
| **TEM-1** | 1.39 | 0.35 | 5.55 | 0.32 |
| **OXA-1 + TEM-1** | 0 |  |  |  |

We then investigated the association between the presence of the OXA-1 and/or TEM-1 genes and susceptibility to piperacillin/tazobactam in *K. pneumoniae* isolates, aggregating data from the three study sites due to the limited number of isolates available for individual site-specific analyses. MIC data were available for 83.6% (56/67) of the isolates, of which 58.9% (33/56) were found to be susceptible to piperacillin/tazobactam. Of the OXA-1-negative isolates, 93.9% (31/33) were susceptible to piperacillin/tazobactam, in contrast to only 8.7% (2/23) of OXA-1-positive isolates.

Similarly, 84.0% (21/25) of TEM-1-negative isolates were susceptible, compared to 38.7% (12/31) of TEM-1-positive isolates. For isolates positive for both OXA-1 and TEM-1, susceptibility was observed in only 5.3% (1/19), compared to 95.2% (20/21) of isolates negative for both genes. Relative risk analysis of non-susceptibility to piperacillin/tazobactam demonstrated highly significant associations with OXA-1 (*P* < 0.001), TEM-1 (*P* = 0.003), and the combination of OXA-1 and TEM-1 (*P* < 0.001) in positive isolates.

We explored the association between the carriage of OXA-1 and/or TEM-1 genes and the presence of other resistance genes, focusing on those associated with resistance to aminoglycosides, fluoroquinolones, trimethoprim/sulfamethoxazole, and tetracycline. Specifically, we examined genes conferring resistance to the following drug classes: aac(3)-IId (aminoglycosides), aac(3)-IIe (aminoglycosides), aac(6’)-Ib-cr5 (aminoglycosides and fluoroquinolones), aadA5 (aminoglycosides), aph(3’’)-Ib (aminoglycosides), aph(6)-Id (aminoglycosides), dfrA17 (trimethoprim), and tet(A) (tetracycline). Figure 2 illustrates the frequency of each gene combination in OXA-1-and/or TEM-1-positive variants. Interestingly, the aac(3)-IId gene was absent in OXA-1-positive variants. Additionally, 98.6% (72/73) of OXA-1-positive variants carried the aac(6’)-Ib-cr5 gene, compared to only 0.72% (2/276) of OXA-1-negative variants. When we analyzed the relative likelihood of OXA-1 being present in relation to other resistance genes, OXA-1-positive variants showed a significant positive association with most of the tested resistance genes; however aph(6)-ld and aph(3”)-lb were significantly less likely to be present when OXA-1 was present. TEM-1-positive variants were significantly associated with the presence of aac(3)-lld, aph(3”)-lb, and aph(6)-ld and the absence of aac(6’)-lb-cr5 and aac(3)-lle (Table 3).

**Table 3.** Data showing relative likelihood of OXA-1 or TEM-1 being present in relation to the presence of other determinant of resistance genes.

| All *E. coli* isolates. Risk of presence with OXA-1 |  |  |  |  |
| --- | --- | --- | --- | --- |
| Resistance gene | RR | 95% lower CI | 95% upper CI | *P* |
| ***aac(3)-Ile*** | 9.33 | 5.43 | 16.03 | **<0.001** |
| ***aac(6')-Ib-cr5*** | 136.11 | 34.2 | 541.65 | **<0.001** |
| ***aadA5*** | 1.37 | 1.04 | 1.8 | **0.011862** |
| ***aph(3'')-Ib*** | 0.58 | 0.37 | 0.92 | **<0.001** |
| ***aph(6)-Id*** | 0.46 | 0.28 | 0.78 | **0.001722** |
| ***dfrA17*** | 1.38 | 1.06 | 1.79 | **0.007917** |
| **tet(A)** | 1.58 | 1.25 | 1.99 | **<0.001** |
| **All *E. coli* isolates. Risk of presence with TEM-1** |  |  |  |  |
| **Resistance gene** | **RR** | **95%**<br>**lower CI** | **95%**<br>**upper CI** | **P** |
| **aac(3)-Ild** | 15.85 | 5.8 | 43.31 | **<0.001** |
| **aac(3)-Ile** | 0.25 | 0.11 | 0.53 | **0.000172** |
| **aac(6')-Ib-cr5** | 0.4 | 0.24 | 0.68 | **<0.001** |
| **aadA5** | 1.17 | 0.91 | 1.51 | 0.117949 |
| **aph(3'')-Ib** | 1.64 | 1.23 | 2.18 | **<0.001** |
| **aph(6)-Id** | 1.56 | 1.17 | 2.08 | **0.001244** |
| **dfrA17** | 1.14 | 0.89 | 1.46 | 0.154633 |
| **tet(A)** | 0.92 | 0.71 | 1.17 | 0.242268 |

**Figure 2.**
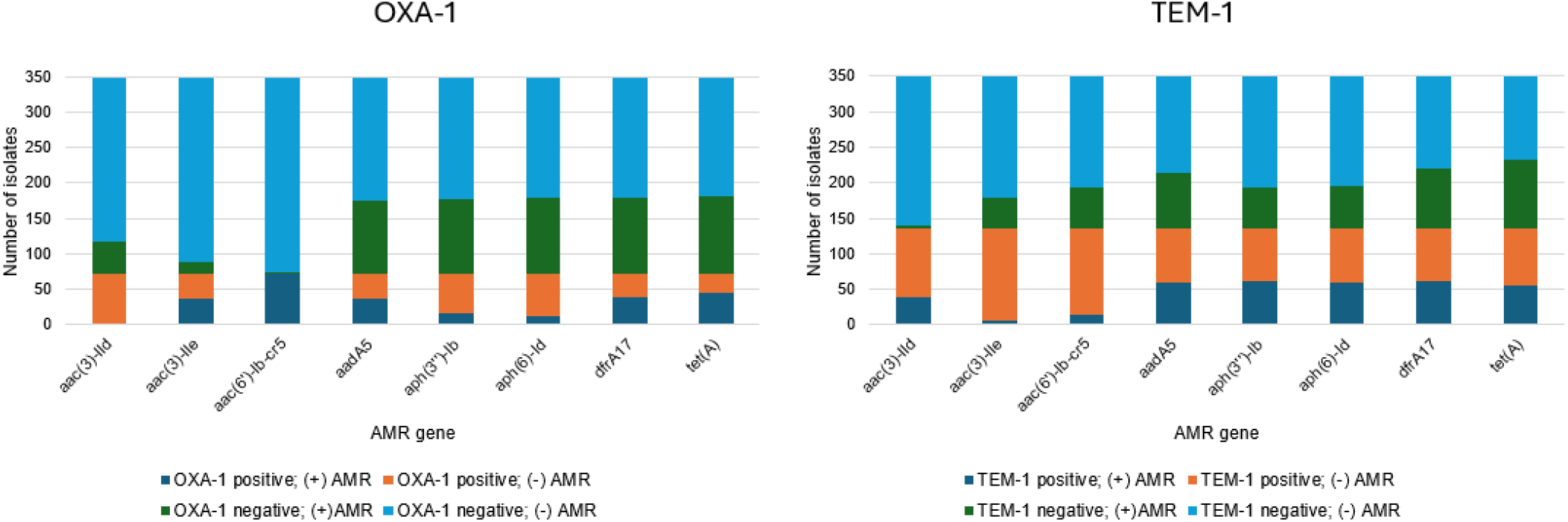
Breakdown of isolates with different combinations of antimicrobial resistance (AMR) genes in *E. coli* isolates. For example, when the aac(6’)-lb-cr5 AMR gene is present, the OXA-1 gene is always present and the aac(6’)-lb-cr5 AMR gene is not present if OXA-1 is not present. On the other hand, when TEM-1 is present the AMR gene aac(3)-lle is rarely present, but is TEM-1 is absent the AMR gene is much more likely to be present

Lastly, we assessed the association between gentamicin susceptibility and the presence of specific resistance genes [aac(3)-IId, aac(3)-IIe, aac(6’)-Ib-cr5, and aph(6)-Id] in OXA-1-or TEM-1 variants. Gentamicin susceptibility data were available for 97.4% (340/349) of the *E. coli* isolates. For the OXA-1 gene no significant association with susceptibility to gentamicin was found (Table 4A). In contrast, the analysis for the TEM-1 gene revealed a significant association with gentamicin susceptibility (Table 4A).

**Table 4A.**
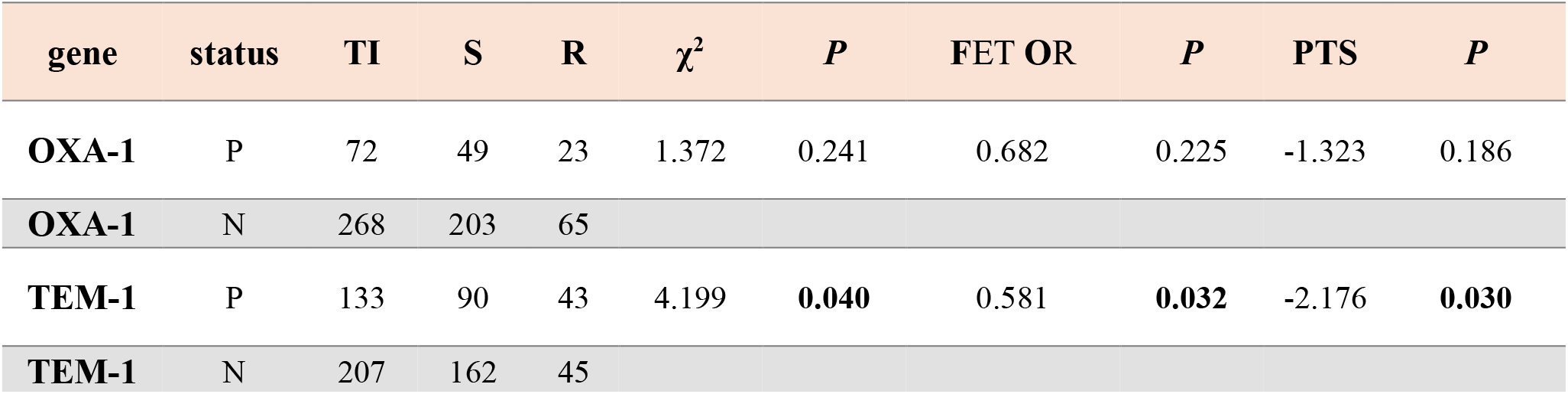
Data showing association of gentamicin susceptibility and the presence of OXA-1 or TEM variants. P – positive; N-negative; TI -Total Isolates; S – susceptible; R – resistant; FET OR - Fisher’s Exact Test (FET) Odds Ratio; PTS - Proportion Test Statistic. Statistically significant values shown in bold.

**Table 4B.**
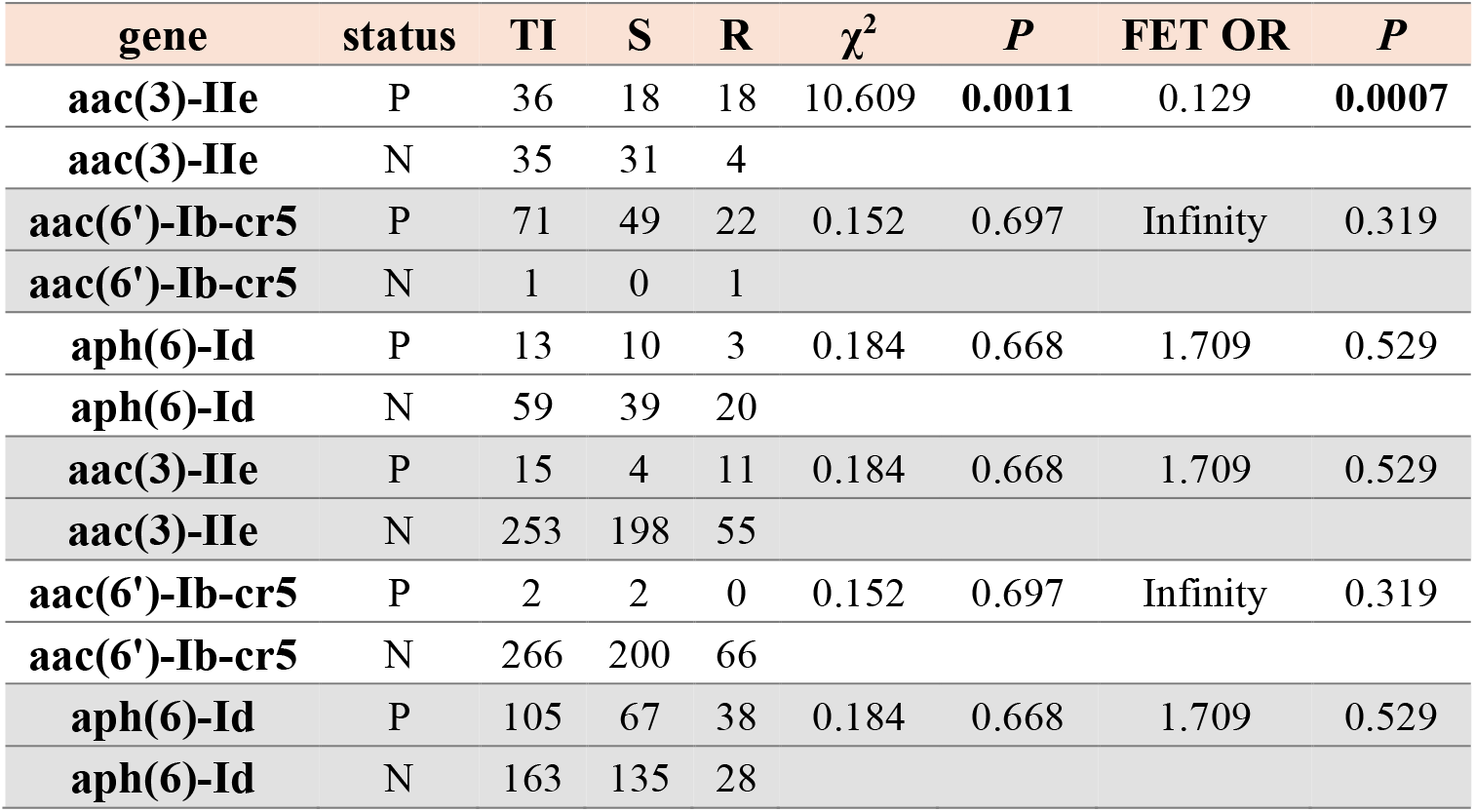
Data showing association of gentamicin susceptibility and the presence of the different genes in OXA-1 variant isolates. Statistically significant values shown in bold.

| gene | status | TI | S | R | $\chi^2$ | P | FET OR | P |
| --- | --- | --- | --- | --- | --- | --- | --- | --- |
| **aac(3)-IIe** | P | 36 | 18 | 18 | 10.609 | **0.0011** | 0.129 | **0.0007** |
| **aac(3)-IIe** | N | 35 | 31 | 4 |  |  |  |  |
| **aac(6')-Ib-cr5** | P | 71 | 49 | 22 | 0.152 | 0.697 | Infinity | 0.319 |
| **aac(6')-Ib-cr5** | N | 1 | 0 | 1 |  |  |  |  |
| **aph(6)-Id** | P | 13 | 10 | 3 | 0.184 | 0.668 | 1.709 | 0.529 |
| **aph(6)-Id** | N | 59 | 39 | 20 |  |  |  |  |
| **aac(3)-IIe** | P | 15 | 4 | 11 | 0.184 | 0.668 | 1.709 | 0.529 |
| **aac(3)-IIe** | N | 253 | 198 | 55 |  |  |  |  |
| **aac(6')-Ib-cr5** | P | 2 | 2 | 0 | 0.152 | 0.697 | Infinity | 0.319 |
| **aac(6')-Ib-cr5** | N | 266 | 200 | 66 |  |  |  |  |
| **aph(6)-Id** | P | 105 | 67 | 38 | 0.184 | 0.668 | 1.709 | 0.529 |
| **aph(6)-Id** | N | 163 | 135 | 28 |  |  |  |  |

**Table 4C.** Data showing association of gentamicin susceptibility and the presence of the different genes in TEM-1 variant isolates. Statistically significant values shown in bold. Same abbreviations as those indicated on Table 4A.

| gene | Status | TI | S | R | chi-square | p-value | FET OR | p-value |
| --- | --- | --- | --- | --- | --- | --- | --- | --- |
| **aac(3)-IId** | P | 39 | 8 | 31 | 10.609 | **0.0011** | 0.129 | **0.0007** |
| **aac(3)-IId** | N | 94 | 81 | 13 |  |  |  |  |
| **aac(3)-IIe** | P | 7 | 3 | 4 | 0.184 | 0.668 | 1.709 | 0.529 |
| **aac(3)-IIe** | N | 126 | 86 | 40 |  |  |  |  |
| **aac(6')-Ib-cr5** | P | 15 | 14 | 1 | 0.152 | 0.697 | Infinity | 0.319 |
| **aac(6')-Ib-cr5** | N | 118 | 75 | 43 |  |  |  |  |
| **aph(6)-Id** | P | 59 | 27 | 32 | 0.184 | 0.668 | 1.709 | 0.529 |
| **aph(6)-Id** | N | 74 | 62 | 12 |  |  |  |  |
| **aac(3)-IId** | P | 4 | 2 | 2 | 10.609 | 0.0011 | 0.129 | 0.0007 |
| **aac(3)-IId** | N | 203 | 160 | 43 |  |  |  |  |
| **aac(3)-IIe** | P | 44 | 19 | 25 | 0.184 | 0.668 | 1.709 | 0.529 |
| **aac(3)-IIe** | N | 163 | 143 | 20 |  |  |  |  |
| **aac(6')-Ib-cr5** | P | 58 | 37 | 21 | 0.152 | 0.697 | Infinity | 0.319 |
| **aac(6')-Ib-cr5** | N | 149 | 125 | 24 |  |  |  |  |
| **aph(6)-Id** | P | 59 | 50 | 9 | 0.184 | 0.668 | 1.709 | 0.529 |
| **aph(6)-Id** | N | 148 | 112 | 36 |  |  |  |  |

We evaluated the association of aac(3)-IIe, aac(6’)-Ib-cr5, aph(3’’)-Ib, and aph(6)-Id genes with gentamicin susceptibility in OXA-1 variants. A significant association between the presence of the aac(3)-IIe gene and susceptibility to gentamicin was found but no significant associations were observed for the aac(6’)-Ib-cr5 or aph(6)-Id genes (Table 4B).

We evaluated the association of aac(3)-IId, aac(3)-IIe, aac(6’)-Ib-cr5, and aph(6)-Id genes with gentamicin susceptibility in TEM-1 variants and found a significant association only for the presence of the aac(3)-IId gene. No significant associations were identified for the other genes analyzed (Table 4C).

## DISCUSSION

This multi-center retrospective study investigated the prevalence of OXA-1 and TEM-1 genes in ESBL-producing *E. coli* and *K. pneumoniae* and their association with susceptibility to piperacillin/tazobactam. The study was conducted across three geographically distinct institutions: TAMC in Hawaii, MAMC in Washington, and BAMC in Southern Texas. This geographic diversity is crucial as it provides insights into regional variations in the prevalence of resistance genes and antibiotic susceptibility patterns. Our findings revealed significant differences in the distribution of resistance genes among the three institutions. For instance, the prevalence of the CTX-M-15 variant was highest at BAMC and lowest at TAMC, while the CTX-M-27 variant was most prevalent at TAMC. These variations underscore the importance of local epidemiological data in guiding empirical antibiotic therapy. The significant association between the OXA-1 gene and reduced susceptibility to piperacillin/tazobactam, particularly at TAMC and MAMC, highlights the need for region-specific treatment strategies. In contrast, no significant association was observed between the TEM-1 gene and susceptibility to piperacillin/tazobactam. Additionally, we found that the presence of the aac(3)-IIe gene was significantly associated with gentamicin resistance in OXA-1 positive variants, while the aac(3)-IId gene was significantly associated with gentamicin resistance in TEM-1 positive variants.

The study emphasizes the necessity of comprehensive genetic profiling and local epidemiological surveillance to inform antibiotic stewardship programs and improve treatment outcomes for patients with ESBL-producing infections. By understanding the regional differences in resistance mechanisms, healthcare providers can tailor antibiotic therapies more effectively, reducing the risk of therapeutic failure and the spread of resistant pathogens.

The emergence of ESBL-producing Gram-negative pathogens has posed significant challenges in clinical settings, particularly in the treatment of infections caused by *E. coli* and *K. pneumoniae*. Our study highlights the importance of understanding the genetic factors contributing to antibiotic resistance, as these can significantly impact treatment outcomes. The significant association between the OXA-1 gene and reduced susceptibility to piperacillin/tazobactam underscores the need for careful consideration of this gene when selecting treatment options for ESBL-producing infections.

The findings from our study are consistent with previous research that has demonstrated the role of OXA-1 in conferring resistance to penicillin/inhibitor combinations. Livermore et al. (2019) reported that the presence of OXA-1 was strongly associated with reduced susceptibility to piperacillin/tazobactam and amoxicillin/clavulanate in ESBL-producing *E. coli* isolates. Similarly, Walkty et al. (2022) found that OXA-1 was associated with elevated piperacillin/tazobactam MIC values among ESBL-producing *E. coli* clinical isolates. These studies, along with our findings, highlight the critical role of OXA-1 in resistance mechanisms and the potential for therapeutic failure when using penicillin/inhibitor combinations in the presence of this gene.

In contrast, our study did not find a significant association between the TEM-1 gene and susceptibility to piperacillin/tazobactam. This is in line with previous studies that have shown variable results regarding the impact of TEM-1 on resistance to penicillin/inhibitor combinations. For instance, Rodríguez-Villodres et al. (2018) demonstrated that piperacillin/tazobactam was less effective against ESBL *E. coli* carrying blaTEM due to increased copy numbers and transcription levels of the gene.

However, other studies have not found a significant impact of TEM-1 on resistance, suggesting that additional factors may influence the efficacy of penicillin/inhibitor combinations in the presence of TEM-1.

Our study also explored the association between the presence of additional resistance genes and susceptibility to gentamicin. The significant association between the aac(3)-IIe gene and gentamicin resistance in OXA-1 positive variants, as well as the association between the aac(3)-IId gene and gentamicin resistance in TEM-1 positive variants, underscores the complexity of resistance mechanisms in ESBL-producing pathogens. These findings suggest that the presence of specific aminoglycoside resistance genes can further complicate treatment outcomes and highlight the need for comprehensive genetic profiling of clinical isolates to guide antibiotic therapy.

The multi-center nature of our study provides valuable insights into the prevalence and distribution of resistance genes across different geographic regions and healthcare settings. The significant differences in the distribution of CTX-M-15, CTX-M-27, and CTX-M-14 variants among the three institutions underscore the importance of local epidemiological data in guiding empirical treatment decisions. The higher prevalence of CTX-M-15 in BAMC, for example, suggests that clinicians in this region may need to be particularly vigilant when selecting antibiotics for ESBL-producing infections.

Our study has several limitations that should be considered when interpreting the findings. The retrospective design may introduce selection bias, and the reliance on MIC data from routine clinical testing may not capture the full spectrum of resistance mechanisms. Additionally, the relatively small sample size for some gene variants may limit the generalizability of the findings. Future studies should aim to include larger sample sizes and prospective data collection to validate and expand upon our results.

In conclusion, our study highlights the critical role of the OXA-1 gene in conferring resistance to piperacillin/tazobactam in ESBL-producing *E. coli* and *K. pneumoniae*. The significant association between specific aminoglycoside resistance genes and gentamicin resistance further underscores the complexity of resistance mechanisms in these pathogens. Our findings emphasize the need for comprehensive genetic profiling and local epidemiological data to guide antibiotic therapy and improve treatment outcomes for patients with ESBL-producing infections.

## MATERIALS AND METHODS

Bacterial isolates

A total of 416 ESBL-producing bacterial isolates from urine or blood specimens submitted to the clinical laboratories for testing in the three MTFs within the DHA as routine patient standard-of-care (SOC) submitted between January 2022 through December 2023 were retrospectively analyzed in this study. The three MTFs are Tripler Army Medical Center (TAMC) in Honolulu, Hawaii; Madigan Army Medical Center (MAMC) located on Joint Base Lewis-McChord, Washington; and Brooke Army Medical Center, located on Joint Base San Antonio-Fort Sam Houston, Texas. These institutions serve active duty service members, their families, military retirees and their families, veterans, and residents of the areas where they’re located, with TAMC serving the Indo-Pacific region, MAMC the Northwestern US region, and BAMC the South Central region of the US.

### Antimicrobial Susceptibility Testing (AST)

Bacterial identification and AST were performed as part of SOC from each patient using VITEK MS and Vitek 2 (bioMérieux, NC, USA) systems. The minimal inhibitory concentration (MIC) were interpreted per Clinical and Laboratory Standards Institute (CLSI) guidelines. ESBL-producing organisms were determined based on Vitek 2 AST algorithm, which included non-susceptibility to ceftriaxone.

### Whole Genome Sequencing and Data Analysis

All ESBL-producing isolates from these and other MTFs are routinely submitted to the Multidrug-resistant organism Repository and Surveillance Network (MRSN) at Walter Reed Army Institute of Research (WRAIR), Silver Spring, MD, for whole genome sequencing (WGS) on the Illumina MiSeq or NextSeq benchtop sequencer (Illumina, Inc., San Diego, CA). Isolation, DNA extraction and library preparation were performed as previously described (13,14). Genome sequencing was performed using either Illumina MiSeq or NextSeq 500 with MiSeq Reagent Kit v3 (2 x 300bp) or NextSeq Reagent Kit 500/550 v2 (2 x 150bp) (Illumina, San Diego, CA, USA). bbduk v38.96 (https://github.com/BioInfoTools/BBMap/blob/master/sh/bbduk2.sh) was used to remove barcode and adapter sequence as well as to perform quality trimming. Kraken2 v2.1.2 (https://github.com/DerrickWood/kraken2) was used initial taxonomic assignment and to screen for contamination. De novo draft genome assemblies were produced using shovill v1.1.0 (https://github.com/tseemann/shovill) with coverage estimates generated using bbmap v38.96. Minimum thresholds for contig size and coverage were set at 200 bp and 49.5+, respectively. Antibiotic resistance genes were identified using ABRicate (https://github.com/tseemann/abricate, v.0.8.13), using the ResFinder database and additional searches using CARD to identify all resistance genes.

### Susceptibility Testing

Susceptibility testing for piperacillin/tazobactam, and gentamicin was performed using the broth microdilution method according to the CLSI guidelines. The minimum inhibitory concentrations (MICs) were determined, and isolates were classified as susceptible or resistant based on CLSI breakpoints.

### Statistical Analysis

Statistical analyses were performed to assess the association between the presence of resistance genes and susceptibility to piperacillin/tazobactam and gentamicin. The chi-square test for independence was used to determine whether there was a significant association between the presence of specific resistance genes and susceptibility to antibiotics. Fisher’s Exact Test was used as an alternative to the chi-square test for small sample sizes or when the assumptions of the chi-square test were not met. The proportion test was used to compare the proportions of susceptible and resistant isolates between different groups. The test statistic and p-values were calculated to determine if there were significant differences in the proportions. Relative risk was calculated to determine the probability of resistance to piperacillin/tazobactam given the presence of certain genes. Relative risk was also calculated for the probability of co-occurrence of antimicrobial resistance markers. All statistical analyses calculations were performed assessing significance at a *P* value equal to 0.05.

### Data Presentation

The results of the statistical analyses were summarized in tables, including the total number of isolates, the number of susceptible and resistant isolates, chi-square test statistics, p-values, Fisher’s Exact Test odds ratios, and p-values for each gene and antibiotic combination.

### Software

All statistical analyses were performed using R software (version 4.0.3) and the stats and exact2×2 packages for chi-square and Fisher’s Exact Tests, respectively. The prop.test function was used for the proportion test.

## DISCALAIMER, ETHICAL STATEMENT AND ACKNOWLEDGEMENT

The views expressed in this study are those of the authors and do not necessarily reflect the official policy or position of the Defense Health Agency, Department of Defense, nor the U.S. Government. This work was prepared as part of official duties. Title 17, U.S.C., Section 105 provides that copyright protection under this title is not available for any work of the U.S. Government. Title 17, U.S.C., Section 101 defines a U.S. Government work as a work prepared by a military service member or employee of the U.S. Government as part of that person’s official duties.

### Ethical statement

The institution’s ethical review board and the human research protection office at the Defense Health Agency and Tripler Army Medical Center reviewed and approved this study. We would like to express our heartfelt gratitude to the laboratory professionals in the microbiology laboratories at Tripler Army Medical Center and across the MTFs. Often working behind the scenes, their expertise, unwavering professionalism, and meticulous attention to detail are instrumental in ensuring the safety and well-being of our patients. Their dedication is a cornerstone of quality care, and we deeply appreciate the critical role they play in supporting our mission. Important as well, we acknowledge the staff at the Walter Reed Army Institute of Research - Multidrug-Resistant Organism Repository and Surveillance Network (WRAIR/MRSN) for whole-genome sequencing of the isolates in this study.

